# Decoding brain basis of laughter and crying in natural scenes

**DOI:** 10.1101/2022.10.11.511708

**Authors:** Lauri Nummenmaa, Tuulia Malèn, Sanaz Nazari-Farsani, Kerttu Seppälä, Lihua Sun, Henry K. Karlsson, Matthew Hudson, Jussi Hirvonen, Mikko Sams, Sophie Scott, Vesa Putkinen

## Abstract

Laughter and crying are universal signals of prosociality and distress, respectively. Here we investigated the functional brain basis of perceiving laughter and crying using naturalistic functional magnetic resonance imaging (fMRI) approach. We measured haemodynamic brain activity evoked by laughter and crying in three experiments with 100 subjects in each. The subjects i) viewed a 20-minute medley of short video clips, and ii) 30 minutes of a full-length feature film, and iii) listened to 15 minutes of a radio play that all contained bursts of laughter and crying. Intensity of laughing and crying in the videos and radio play was annotated by independent observes, and the resulting time series were used to predict hemodynamic activity to laughter and crying episodes. Multivariate pattern analysis (MVPA) was used to test for regional selectivity in laughter and crying evoked activations. Laughter induced widespread activity in ventral visual cortex and superior and middle temporal and motor cortices. Crying activated thalamus, cingulate cortex along the anterior-posterior axis, insula and orbitofrontal cortex. Both laughter and crying could be decoded accurately (66-77% depending on the experiment) from the BOLD signal, and the voxels contributing most significantly to classification were in superior temporal cortex. These results suggest that perceiving laughter and crying engage distinct neural networks, whose activity suppresses each other to manage appropriate behavioral responses to others’ bonding and distress signals.

**Significance statement:** Laughter and crying are universal signals of prosociality and distress, respectively. They occur in complex, dynamic social settings with variable and dynamically evolving time courses. Here we used functional magnetic resonance imaging experiments and statistical pattern recognition for disentangling the neural systems that encode laughter and crying signals from dynamic and highly naturalistic scenes. These results show that separable neural circuits are engaged in processing distinct types of social attachment cues, and that pattern recognition during dynamic scene perception allows reliable separation of laughter and crying evoked activation patterns. Coordinated activity of these networks allows managing appropriate behavioral responses to others’ bonding and distress signals.

## Introduction

Humans have an urgent need to feel belonging to groups and use a multitude of expressions for signifying this. Laughter is a universally recognized positive social expression. It occurs frequently in human social interactions (Sauter et al., 2010; Scott et al., 2015) but is also common among nonhuman primates (Preuschoft, 1992; Ross et al., 2009) and rodents (Panksepp and Burgdorf, 2003). Macaques and chimpanzees use a quiet smile-like gesture to appease aggressive conspecifics, whereas relaxed open-mouth vocalizations are associated with both play behavior and pair formation (Preuschoft, 1992; Waller and Dunbar, 2005). Similarly, humans use quiet smiles for signaling social approval and openness to social interaction (Calvo et al., 2012; Calvo and Nummenmaa, 2015), whereas laughter is used directly for promoting social bonding (Dunbar, 2012; Scott et al., 2015). Functional and acoustic properties these play signals in humans resemble those of numerous other animals, most notably other great apes, suggesting shared evolutionary origin (Winkler and Bryant, 2021). Human molecular imaging studies have shown that the bonding function of laugher is governed by the endogenous opioid system (Manninen et al., 2017; Sun et al., 2022) that modulates both pleasurable and calm sensations (Nummenmaa and Tuominen, 2018; Kantonen et al., 2020). Crying is also used for signaling the need for social contact, but unlike laughter it is evoked when social losses or social distancing is experienced. Crying engages the separation distress circuit in the mammalian brain that consequently modulates approach behavior and social contact seeking (Panksepp, 2003). Due to the centrality attachment in mental health and well-being and, it is critical to understand the functional systems processing of these distinct types of social attachment signals.

There is a surprising paucity of data on the neurobiology of the social bonding circuits engaged by laughter and crying. For example, the most widely used sets of static and dynamic human facial emotional expressions (Ekman and Friesen, 1976; Lundqvist et al., 1988; Tottenham et al., 2009; van der Schalk et al., 2011) explicitly exclude laughter and crying from the expression categories. Similarly, the NeuroSynth database for fMRI activation meta-analysis (Yarkoni et al., 2011) does not contain a sufficient number of studies for generating meta-analysis for terms “laughter” or “crying”. The extant literature however shows that laughter generation involves the motor and supplementary motor cortices, and limbic regions inlcuding anterior cingulate cortex, amygdala, nucleus accumbens, and hippocampus. Further modulatory systems include basal ganglia, thalamus, and cerebellum (Talami et al., 2020; Gerbella et al., 2021). Crying, in turn is generated via the interplay between medulla and midbrain structures as well as the hypothalamus, amygdala, insula and prefrontal cortices (Newman, 2007; Bylsma et al., 2019). Hear adult laughter and crying activates partially overlapping regions as laughing and crying, most notably the amygdala, insula, and auditory cortices (Sander and Scheich, 2001; Sander et al., 2003; Sander and Scheich, 2005b; Fecteau et al., 2007). Pattern recognition studies have also found that vocal affect bursts including laughter and crying can be successfully decoded from the brain activity in the auditory and inferior frontal cortices (Kotz et al., 2013; Paquette et al., 2018). However, such focal, effects are in stark contrast with the widespread activation of limbic and paralimbic circuits typically activated during emotional episodes (Kober et al., 2008; Nummenmaa et al., 2012; Nummenmaa et al., 2014).

### The current study

Laughter and crying occur in complex social settings with dynamically evolving time courses. However, all the previous studies on laughter and crying have measured brain responses to isolated acoustic segments. Accordingly, it can be questioned whether these data generalize to the complex and dynamic real-world affiliative behaviour (Adolphs et al., 2016). Here we measured brain responses to laughter and crying in three large-scale (n=100) fMRI experiments using naturalistic audiovisual and auditory stimuli. All experiments yielded reliable and dissociable responses to laughter and crying, and statistical pattern recognition further allowed accurate classification of laughter and crying episodes, with voxels in the auditory cortices contributing most consistently to the classification.

## Methods

Altogether 102 volunteers (51 females, mean age 31 years, range 20–57 years) participated in the study and completed all the three experiements. The exclusion criteria included a history of neurological or psychiatric disorders, alcohol or substance abuse, current use of medication affecting the central nervous system and the standard MRI exclusion criteria. Two additional subjects were scanned but excluded from further analyses because of unusable MRI data due to gradient coil malfunction. All subjects gave an informed, written consent and were compensated for their participation. The ethics board of the Hospital District of Southwest Finland had approved the protocol and the study was conducted in accordance with the Declaration of Helsinki.

### Experimental design

Three experiments were conducted to map the brain basis of perceiving natural laughter and crying and all subjects completed all the experiments. First, to map responses to natural audiovisual laughter we used our previously validated movie viewing paradigm in which the subjects viewed a compilation of 96 movie clips extracted from mainstream English language feature films (mean duration 12.5 s; total duration 20 minutes) containing variable emotional and non-emotional content (for details, see Karjalainen et al., 2017, 2019; Lahnakoski et al., 2012). The movie clips were presented in fixed order without breaks in between and contain no coherent plot structure when viewed after each other. Second, subjects viewed the first 30 minutes of the Finnish feature film *Käsky* (*The Commandment* by Aku Louhimies / Helsinki Filmi, 2008). This allowed testing for generalization of the brain responses to laughter with two independent audiovisual stimulus sets of which one contained a clear narrative structure. Third, to measure responses to naturalistic acoustic laughter only, the subject listened to the first 10 minutes of a radio play *Puhdistu*s (*Purge*, Sofi Oksanen / Radioteatteri, 2011). The film clips were presented via NordicNeuroLab VisualSystem binocular display, sound was delivered binaurally via MRI-compatible headphones (Sensimetrics S14) at a comfortable level adjusted individually for each participant. During the radio play a fixation cross was shown on the screen. Subjects were instructed to attend the stimuli similarly as they were viewing a movie or listening to a podcast, other than that there was no specific task. For all the stimuli used in the three experiments, dynamic ratings with a 4 sec temporal resolution were obtained for the intensity of perceived laughter and crying from a separate sample of subjects (n=6) who did not participate in the fMRI study. The average ratings were subsequently used as regressors in GLM analysis.

### MRI data acquisition

The MRI data were acquired using a Phillips Ingenuity TF PET/MR 3T whole-body scanner. High-resolution (1 mm^3^) structural images were obtained with a T1-weighted sequence (TR 9.8 ms, TE 4.6 ms, flip angle 7°, 250 mm FOV, 256 × 256 reconstruction matrix). A total of 407 functional volumes were acquired with a T2*-weighted echo-planar imaging sequence (TR 2600 ms, TE 30 ms, 75° flip angle, 240 mm FOV, 80 × 80 reconstruction matrix, 62.5 kHz bandwidth, 3.0 mm slice thickness, 45 interleaved slices acquired in ascending order without gaps). Significant gross brain pathology was excluded with T2-weighted images.

### Structural and functional MRI data preprocessing

MRI data were preprocessed using fMRIPprep 1.3.0.2 (Esteban, Markiewicz, et al. 2018). The following preprocessing was performed on the anatomical T1-weighted (T1w) reference image: correction for intensity non-uniformity, skull-stripping, brain surface reconstruction, spatial normalization to the ICBM 152 Nonlinear Asymmetrical template version 2009c (Fonov et al., 2009) using nonlinear registration with antsRegistration (ANTs 2.2.0) and brain tissue segmentation. The following preprocessing was performed on the functional data: co-registration to the T1w reference, slice-time correction, spatial smoothing with a 6mm Gaussian kernel, automatic removal of motion artifacts using ICA-AROMA (Pruim et al., 2015) and resampling the MNI152NLin2009cAsym standard space. Low-frequency drifts were removed with a 240-s-Savitzky–Golay filter (Cukur et al., 2013).

### GLM data analysis

The fMRI data analyzed in SPM12 (Wellcome Trust Center for Imaging, London, UK, (http://www.fil.ion.ucl.ac.uk/spm). To reveal regions activated by laughter and crying, a separate general linear model (GLM) was fitted to the data where the BOLD signal was modelled with the laughter / crying intensity regressors as parametric modulators. Data were analyzed in two ways: First, by modelling all three experiments together (as all subjects completed all the experiment) and second, by modelling each experiment individually to address the generalizability of the effects across the experiments. Contrast images for the main effects of laughter and crying were generated for each participant and subjected to a second-level (random effects) analysis for population-level inference. The statistical threshold was set a p<0.05, FWE corrected.

To visualize the laughter and crying dependent responses, regional effects (betas) for laughter and crying were extracted in bilateral anatomical regions of interest (ROIs) in occipital (inferior occipital and fusiform gyrus), parietal (angular and posterior cingulate gyrus and precuneus), temporal (Heschl’s gyrus, planum temporale, and planum polare), frontal (frontal pole, middle and inferior occipital gyrus), and limbic (nucleus accumbens, amygdala, and thalamus) regions. The betas were then subjected to repeated measures t tests to reveal significantly different regional responses to laughter versus crying.

### Statistical pattern recognition

A between-subject classification of natural laughter and crying was performed in datasets from three experiments including movie clips, feature film, and radio play, separately. For the classification, one label (either laughter or crying) was assigned to each time point of the signal based on the observation and evaluation of 6 raters (see experimental design section for more details). Each dataset was divided into 10 chunks and all time-points with the same class in a chunk were considered as a single event of that class. We checked the distribution of the chunks and confirmed that timepoints with the same label were not interspersed into different chunks; otherwise temporal autocorrelation of adjacent timepoints could result in an artificially increased classification accuracy. The average chunk length was 120 seconds for the movie clips. 198 seconds for the full movie and 83 seconds for the radio play; overall the data contained 20 events for each subject. Chunk-wise GLM with regressors for each class (laughter and crying) was fit to the data resulting in 20 beta weights (2 classes × 10 events per class) as input for the MVPA. The beta weights were then normalized (μ = 0, σ = 1) before the application of MVPA.

An SVM classifier with radial basis function (RBF) kernel was trained to identify the laughter and crying using leave-one-subject-out cross-validation, where the model was trained on the data from all except one subject and tested on the hold-out subject data. We repeated this procedure 100 times so that each subject was once considered as the hold-out subject. This analysis was performed on each experiment’s dataset (movie clips, Finnish feature film, and radio play) using whole-brain images which were previously skull-striped. We performed an ANOVA feature selection to the training set within each cross-validation where 5000 voxels with the highest F-score were selected. We calculated the accuracy of the classifier by computing the proportion of correctly classified events relative to the total number of events. The MVPA analyses were performed in Python using the PyMVPA toolbox (Hanke et al. 2009).

## Results

Figure 2. shows the time series of the laughter and crying bursts in each experiment. We first ran a joint analysis across the experiments to test which brain regions are activated by laughter and crying irrespective of the stimulation type (**Figure 3**). This revealed that both laughter and crying activated auditory cortices and inferior and ventral temporal cortical areas, yet the auditory responses were more clearly bilateral for laughter. Furthermore, laughter was associated with significant activation in the motor and lateral frontal cortex, and deactivations in the anterior, middle, and posterior cingulate cortices, as well as temporo-parietal junction (TPJ). Conversely, crying was associated with increased activation in the posterior cingulate and precuneus, in addition to the medial frontal and thalamic activations. Direct comparison between laughter versus crying revealed that laughter only elicited stronger activations in the left superior temporal cortex, while crying-related responses were stronger in the inferior and ventral occipital regions, throughout the parietal, temporal, and frontal cortices as well as in the thalamus (**Figure 3-4**).

**Figure 1.**
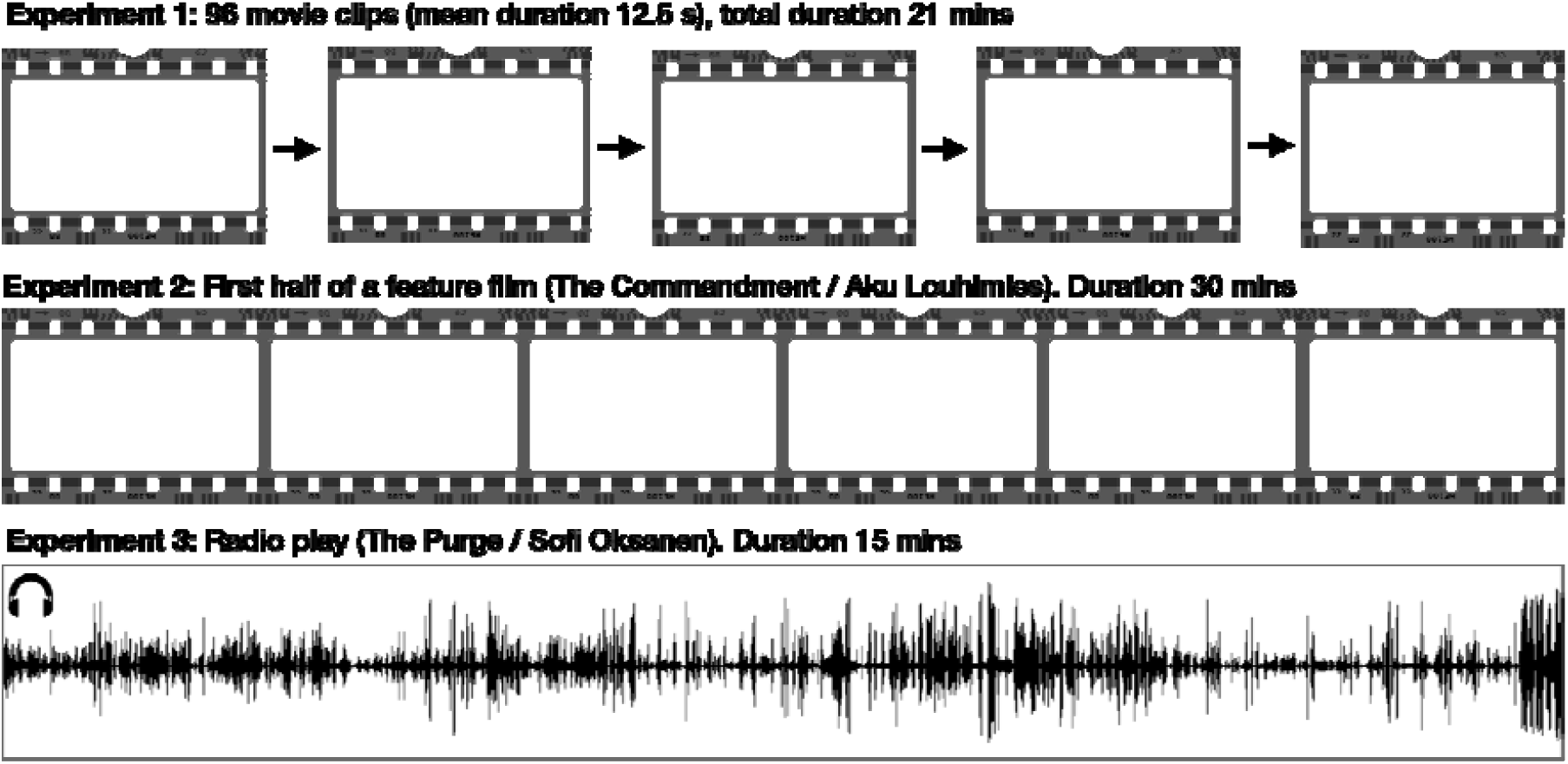
Experimental setup. The subjects viewed a medley of 96 movie clips (mean duration 12.5 s) in a fixed order (Experiment 1), 30 minutes of a feature film (Experiment 2) and listened to 15 minutes of radio play (Experiment 3). Intensity of laughter and crying in the movie clips was annotated at 0.25Hz temporal resolution. Note – sample images have been removed due to bioRxiv policy.

**Figure 2.**
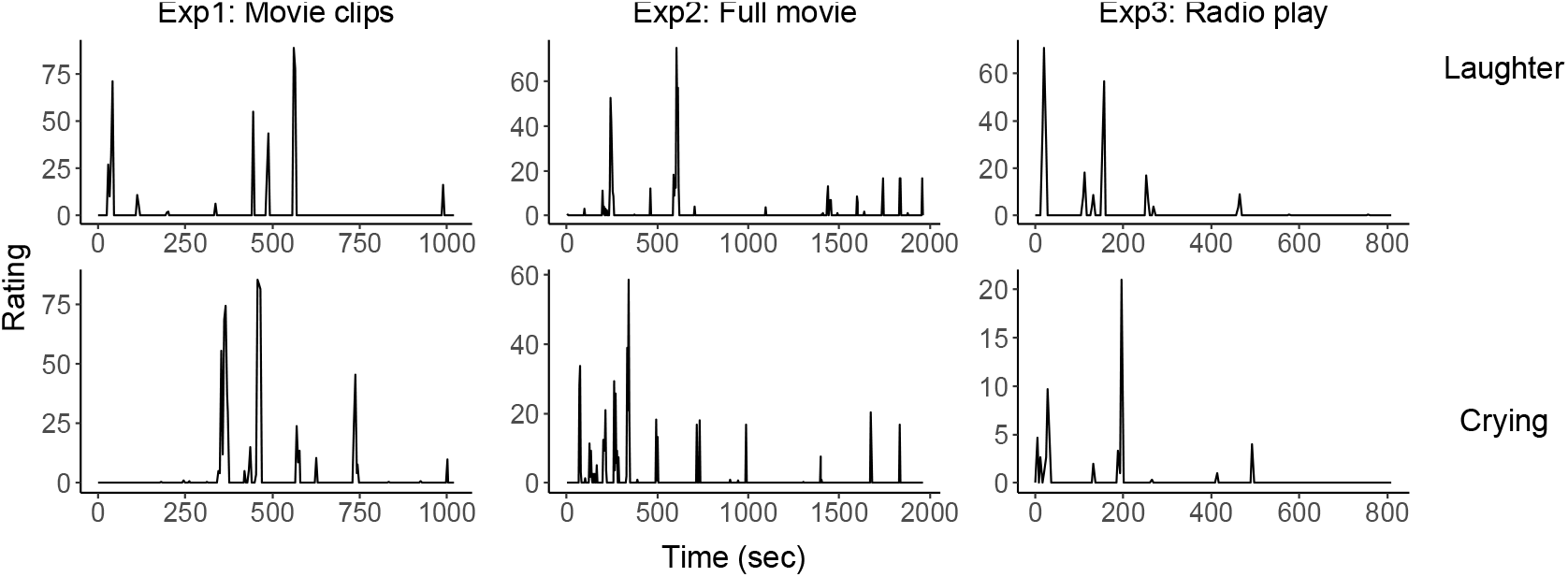
Time series for the laughter and crying regressors in each experiment.

**Figure 3.**
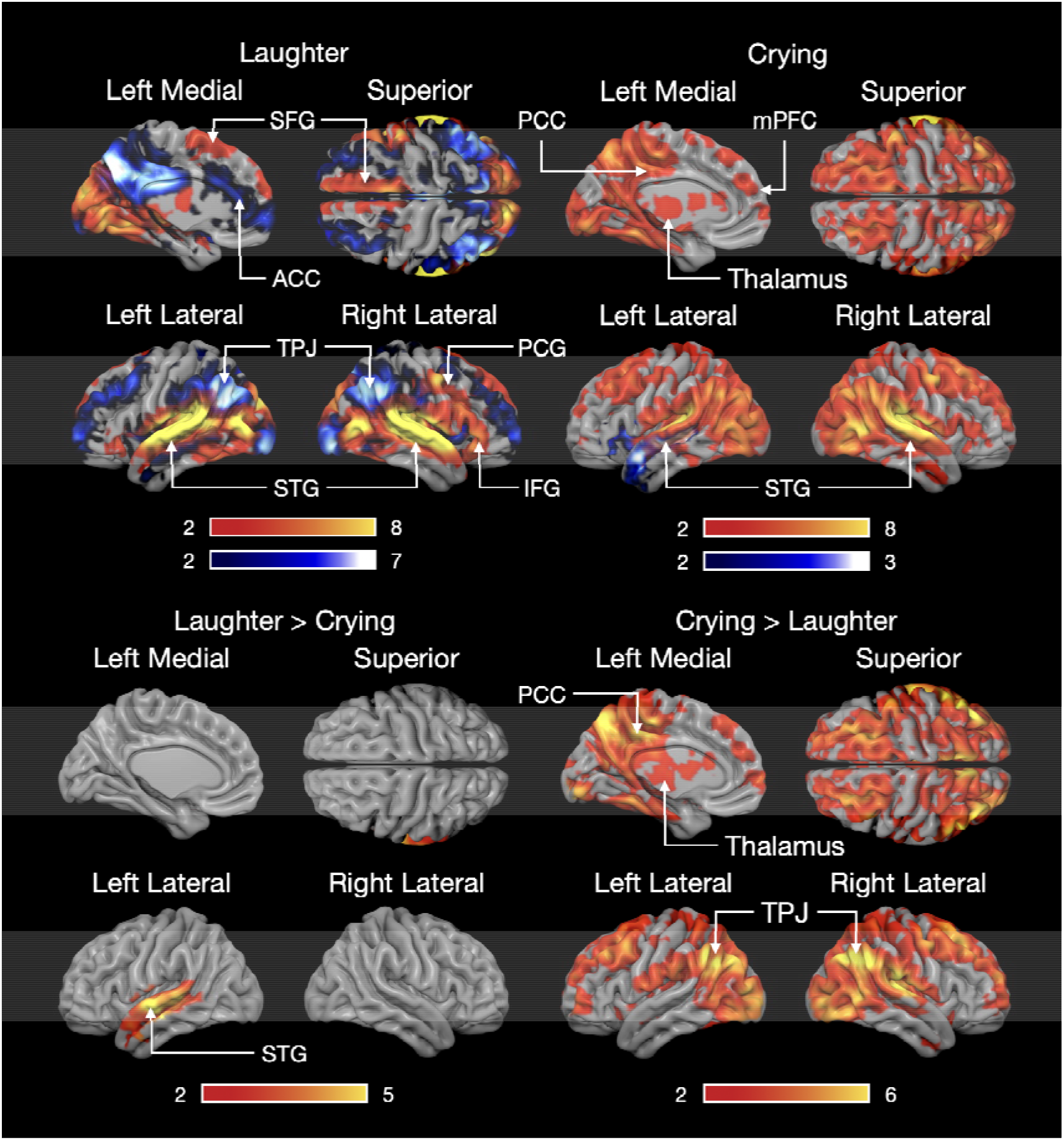
Brain responses to crying and laughter across all experiments. The data are thresholded at p <0.05 FWE corrected. Colourbar shows the t-statistic range. ACC= Anterior Cingulate Cortex, IFG = Inferior Frontal Gyrus, mPFC= Medial prefrontal cortex, PCC= Posterior cingulate, PCG = Precentral Gyrus, STG= Superior Temporal Gyrus, TPJ= Temporo-Parietal Junction.

**Figure 4.**
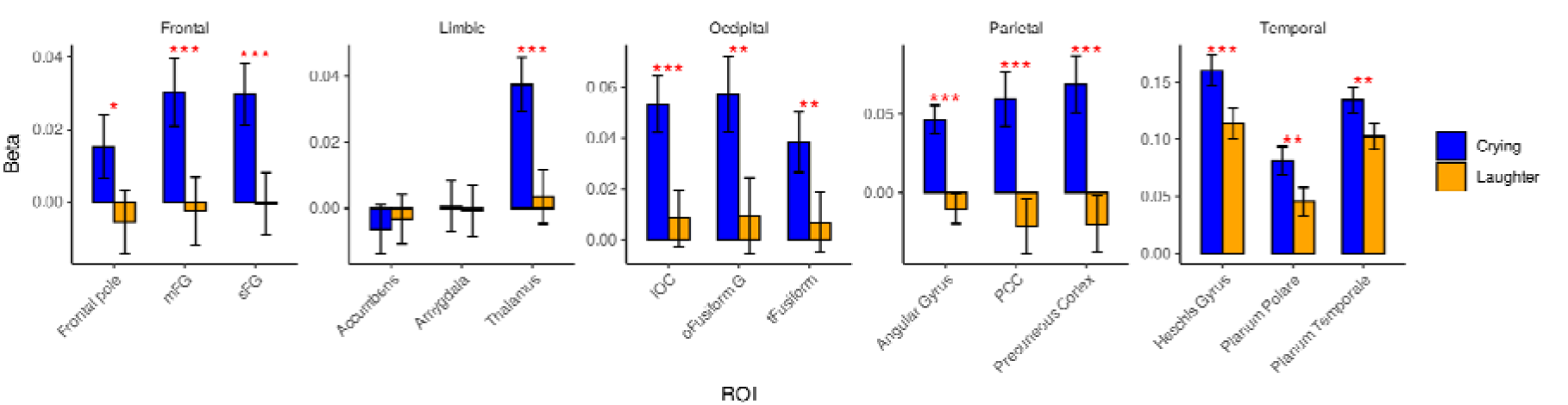
Regional responses (betas) to laughter and crying across all experiments. * = p < 0.05, ** = p < 0.01, *** = p < 0.001

We next tested for the main effect of laughter and crying separately for each experiment. For laughter, this analysis (**Figures S1-S2**) indicated consistent activation of the auditory cortices to laughter in all the experiment. In the audiovisual experiments significant activations were also found in the brainstem, thalamus, V1, ventral temporal and lateral superior temporal cortices, somaotosensory, motor, and lateral frontal cortices. Amygdala activations were most prominent in the experiment with movie clips, whereas striatal activations were strongest in the experiment with the full movie. In the audio only experiment the responses were restricted to the auditory and right lateral frontal cortex. Crying also activated the auditory cortices consistently across experiments. Additional consistent activations were found in the posterior cingulate cortex, somatosensory cortex and parts of the frontal and middle cingulate cortices. In the experiment with movie segments, crying resulted in large-scale deactivations in the primary visual cortex as well as throughout the temporal visual areas.

### Statistical pattern recognition

Because both laughter and crying activated overlapping areas (most notably in the auditory cortex) we next assessed whether laughter and crying nevertheless elicited statistically discernible activation patterns in these and other areas. We ran the pattern recognition for the whole-brain grey matter mask, which yielded classification accuracies that significantly exceeded the permutation-based chance level. Classification accuracies generally exceeded 0.66 for both laughter and crying in all experiments (**Figure 5**), being the most accurate in the experiments with audiovisual stimuli and least accurate in the audio-only experiment. The voxels contributing most significantly to accurate classification were in the auditory cortices in all the experiments, with comparable foci in the superior temporal cortices across all experiments.

**Figure 5.**
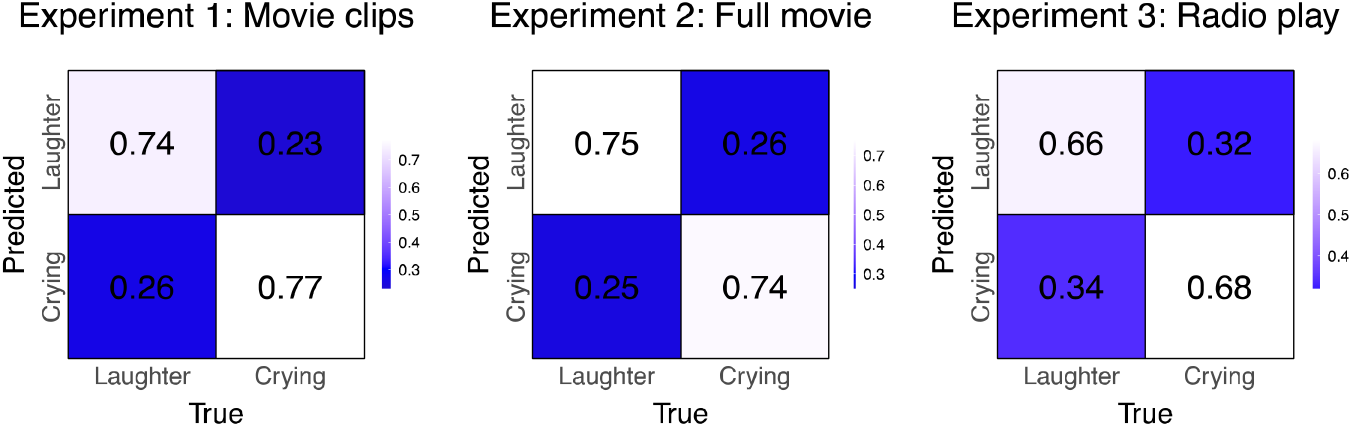
Confusion matrices for decoding laughter and crying in the Experiments 1-3.

**Figure 6.**
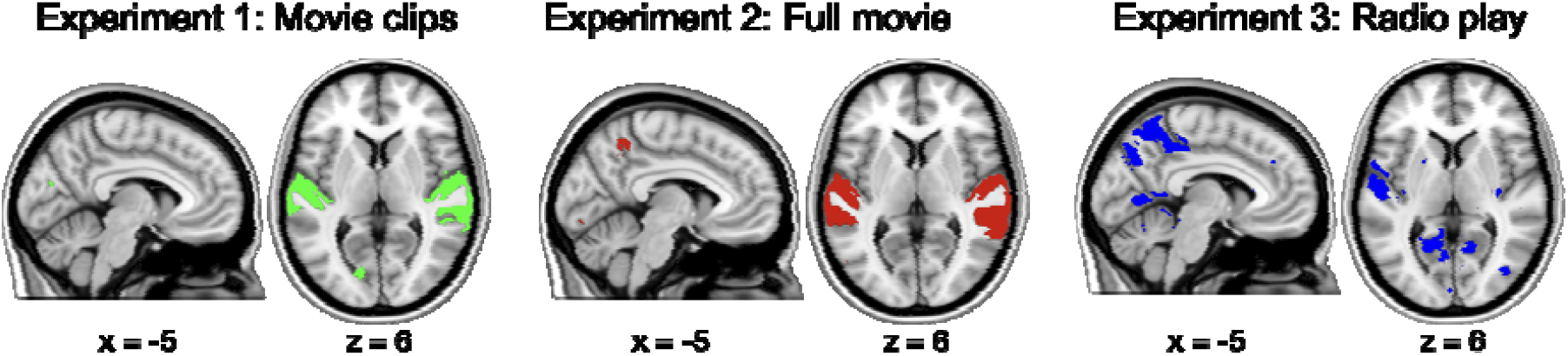
Voxels contributing most significantly to the accurate classification of laughter and crying across the three experiments.

## Discussion

Our main finding was that perceiving laughter and crying from natural scenes engages distinct cortical and subcortical networks. These responses were moderately consistent across the three experiments, although in the acoustic-only experiment they were focused more narrowly on the auditory cortices. The frontoparietal and cingulate regions showed opposite activation patterns with laughter and crying, with decreased activity during laughter episodes and increased activity during crying episodes. While auditory cortices were consistently activated across experiments for laughter and crying sounds, statistical pattern recognition revealed distinct activation patterns for both vocalization types primarily in the auditory cortex. These results show that separable neural circuits are engaged in processing distinct types of social attachment cues, and that pattern recognition during dynamic scene perception allows reliable separation of laughter and crying evoked activation patterns.

### Brain responses to naturalistic laughter and crying

Our data revealed that naturalistic laughter and crying evoked widespread cortical and subcortical activation patters that extend well beyond the auditory cortices. For laugher, most prominent activations were observed in M1, lateral frontal cortex and thalamus. These effects were paralleled by significant deactivations in the anterior, middle, and posterior cingulate cortices as well as temporo-parietal junction (TPJ). The latter set of regions however showed the opposite pattern during crying sounds: These regions were significantly *more* active during the perception of the crying sounds. In addition to these regions, crying robustly activated the posterior cingulate / areas, in addition to the medial frontal and thalamic activations. This is in contrast with prior unimodal studies that have found that laughter and crying evoked activations focus primarily on the auditory cortex, amygdala, and insula (Sander et al., 2003; Wild et al., 2003; Sander and Scheich, 2005a; Sander and Scheich, 2005b; Fecteau et al., 2007).

Grooming-based social bonding imposes constraints on the maximum possible social network size (Dunbar, 1991). Consequently, ecological pressures demanding larger group sizes have led to the evolution of more effective mechanisms for facilitating social bonding (Dunbar, 2008). Laughter is a pleasant prosocial signal that is highly contagious (Scott et al., 2015) and prior studies have indicated that laughter also induces activation of the motor and premotor areas (Lavan et al., 2017). Such “mirroring” of laughter may serve social bonding, as it presumably allows effective spreading of laughter across large crowds (Dunbar, 2012; Manninen et al., 2017), and both behavioral experiments and positron emission tomographic studies indicate that the calming effects of the laughter-evoked endogenous opioid release act as the safety signal promoting subsequent seeking of similar social contacts (Manninen et al., 2017).

However, we also observed significant motor cortex activation for the crying sounds. There is evidence for contagious crying and affect sharing already in infants (Simner, 1971; Geangu et al., 2010) and meta-analysis of functional imaging studies have found that perceiving infant crying activates the dorsal anterior insula, the pre-supplementary motor area and dorsomedial prefrontal cortex and the inferior frontal gyrus, as well as thalamus and cingulate cortices (Witteman et al., 2019). This kind of somatomotor “mirroring” of sadness may promote social behavior by synchronizing the thoughts and feelings across individuals, and fMRI studies using intersubject correlation analyses have indeed found that brain activation in the middle/anterior cingulate cortices becomes increasingly synchronized across individuals during negative emotional states (Nummenmaa et al., 2012; Nummenmaa et al., 2014). The anterior cingulate cortex is a part of the putative separation distress circuit in humans, responding to the perceived physical or affective proximity of conspecificts (Panksepp, 2003). The cingulate cortex acts as a key node of the human saliency network (Bressler and Menon, 2010), it is possible that the crying-evoked cingulate activity reflects the orienting response towards the abrupt yet socially highly relevant distress call. However, the insular cortex is also centrally involved in interoceptive processing and particularly so during emotions (Craig, 2002; Critchley and Garfinkel, 2017); the present fMRI experiments cannot however reveal which of these mutually non-exclusive roles of the insula better explains the data.

### Discrete neural signatures for laughter and crying

Across the three experiments we were consistently able to classify the presence of laughter and crying with above chance level accuracy. Unlike prior studies on vocal affect categorization (Kotz et al., 2013; Paquette et al., 2018), we classified brain signals evoked by unstructured and uncontrolled, dynamic naturalistic stimuli. Despite the complex unstructured stimulus, we nevertheless achieved high classification accuracies particularly in the audiovisual experiments. This shows how pattern recognition can be used for disentangling the specific social perceptual processes that are embedded in high-dimensional and dynamic sensory input. Voxels contributing most significantly to classification of the laughter and crying sounds were localized in the superior temporal cortices. This is in line with previous studies on decoding of auditory affective signals such as vocalizations and music, in which classification can typically be achieved in the auditory cortices (Kotz et al., 2013; Paquette et al., 2018; Putkinen et al., 2021).

Yet, these findings are in stark contrast with patter recognition studies of emotions evoked by e.g. film clips or mental imagery, which consistently suggest discernable and emotions-specific activation patterns in the limbic and paralimbic emotion circuits (Kragel and Labar, 2015; Kragel et al., 2016; Saarimäki et al., 2016; Saarimäki et al., 2018). Despite high statistical power with 100 subjects and long naturalistic experiment, we found no evidence for discernible activation patterns for laughter and crying in the limbic or subcortical regions in general. These findings can likely be reconciled by the fact that vocal expressions of laughter and crying are communicative signals rather than direct readouts of an individual’s emotional state, and the corresponding acoustic-communicative differences are picked up by the multivariate classifier. Thus, it is not unexpected that their processing does not necessarily lead to emotion-specific activation in the subcortical circuits that govern affective processing similarly as for the perception of actual emotion-eliciting episodes (Nummenmaa and Saarimäki, 2017).

## Limitations

Laughter is a complex social signal. Although it is used for signaling social affiliation, it can also be used to signal rejection, and laughter-evoked activations differ depending on the perceived positive versus negative intent (Ethofer et al., 2020). Also, behavioural and neuroimaging studies have revealed that humans are sensitive in spotting genuine versus volitional laughter, which also elicit distinct neural activation patterns (Scott et al., 2015; Lavan et al., 2017). Tractographic studies suggest that emotional and nonemotional (conversational or social) laughter are subserved by partially separable neural networks (Gerbella et al., 2021). We did not specifically annotate for the type of laughter in the video clips and the radio play thus it remains unresolved how specific versus generalizable these effects are across different laughter types. Also, we deliberately used unconstrained naturalistic stimuli and did not define the stimulation a priori, but instead used subject-based annotations for deriving it. However, to overcome the generalizability issue we also pooled data from three experiments (total duration >60 minutes) with 100 subjects in each, and robust GLM activations as well as significantly above-chance level classification accuracy in all experiments. Separate analysis of the three experiments revealed that the laughter and crying evoked activations were not completely consistent across different experiments (**Figure S-1**), highlighting the intrinsic variability in the naturalistic stimulus episodes occurring in the three experiments. Despite variation across exp the across-experiments GLM effects for laughter and crying (**Figure 3**) can be considered as the regions that are most consistently activated while perceiving laughter and crying across variable naturalistic contexts.

## Conclusions

Laughter and crying engage both shared and distinct cortical and subcortical circuits. Although they both trigger robust activation in the auditory cortex, these sensory cortical responses allow reliable encoding of the whether laugher or crying was present in the current audiovisual segment. Activity within the cortical midline network altered between laughter and crying episodes. These results suggest that perceiving laughter and crying engage distinct neural networks, whose activity suppresses each other to manage appropriate behavioral responses to others’ bonding and distress signals.

## Acknowledgements

The study was supported by the Sigrid Juselius Foundation and Academy of Finland (grants numbers 294897 and 332225, to L.N.), the Päivikki and Sakari Sohlberg Foundation (personal grant to T.M.) and the State research funding for expert responsibility area (ERVA) of the Tyks Turku University Hospital (T.M. and L.N.).

