## Supplementary information for "Decoding brain basis of laughter and crying in natural scenes"

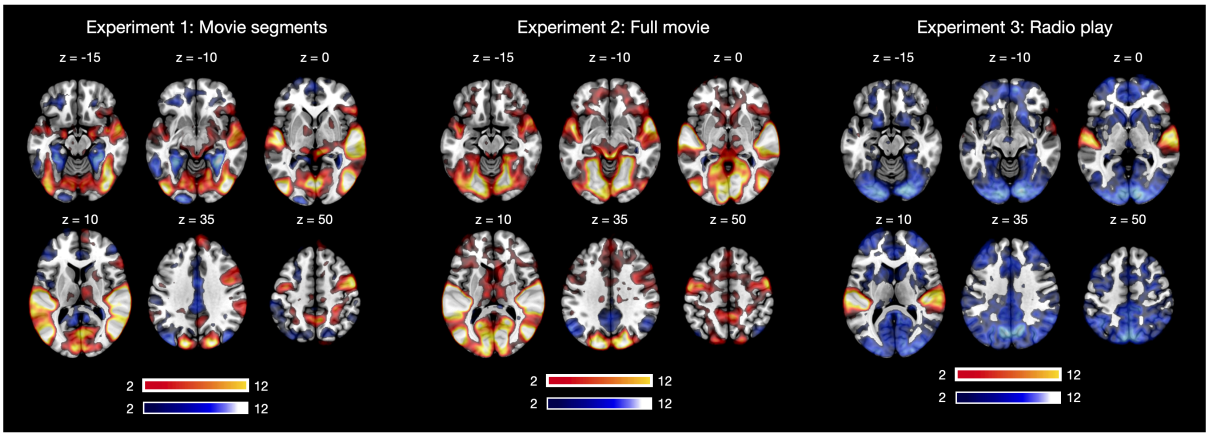


**Figure S-1** Brain responses to laughter in each experiment. The data are thresholded at p <0.05 FWE corrected. Colourbar shows the t-statistic range.

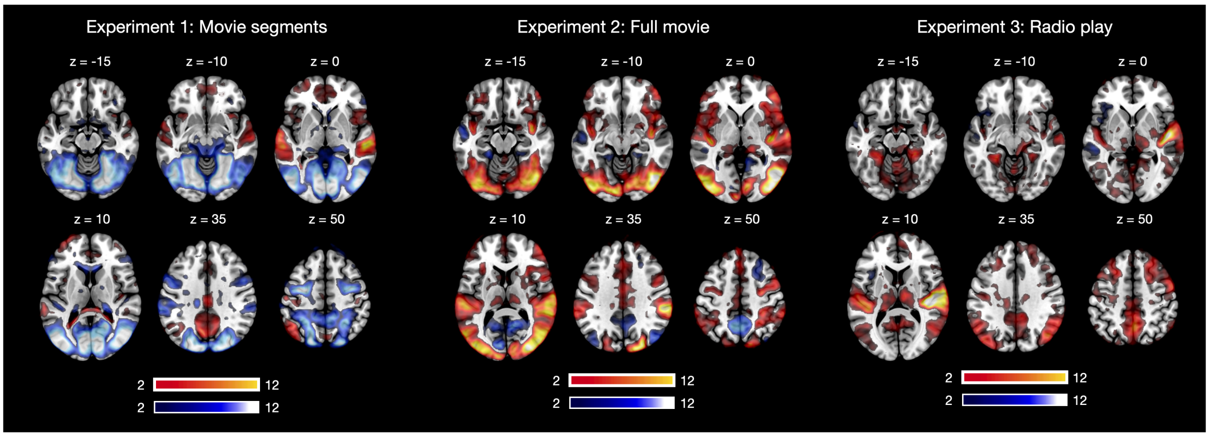


**Figure S-2** Brain responses to crying in each experiment. The data are thresholded at p <0.05 FWE corrected. Colourbar shows the t-statistic range.
